# Evaluating the performance of selection scans to detect selective sweeps in domestic dogs

**DOI:** 10.1101/028647

**Authors:** Florencia Schlamp, Julian van der Made, Rebecca Stambler, Lewis Chesebrough, Adam R. Boyko, Philipp W. Messer

## Abstract

Selective breeding of dogs has resulted in repeated artificial selection on breed-specific morphological phenotypes. A number of quantitative trait loci associated with these phenotypes have been identified in genetic mapping studies. We analyzed the population genomic signatures observed around the causal mutations for 12 of these loci in 25 dog breeds, for which we genotyped 25 individuals in each breed. By measuring the population frequencies of the causal mutations in each breed, we identified those breeds in which specific mutations most likely experienced positive selection. These instances were then used as positive controls for assessing the performance of popular statistics to detect selection from population genomic data. We found that artificial selection during dog domestication has left characteristic signatures in the haplotype and nucleotide polymorphism patterns around selected loci that can be detected in the genotype data from a single population sample. However, the sensitivity and accuracy at which such signatures were detected varied widely between loci, the particular statistic used, and the choice of analysis parameters. We observed examples of both hard and soft selective sweeps and detected strong selective events that removed genetic diversity almost entirely over regions >10 Mbp. Our study demonstrates the power and limitations of selection scans in populations with high levels of linkage disequilibrium due to severe founder effects and recent population bottlenecks.

## INTRODUCTION

Identifying the molecular targets on which positive selection has acted constitutes one of the key challenges for modern population genetics. Ideally, positive selection is inferred directly from the frequency changes of selected alleles in a population over time (Malaspinas *et al*. 2012; Foll *et al*. 2014; Bank *et al*. 2014). However, such approaches require data on historic allele frequencies, otherwise they remain limited to situations of particularly rapid evolution that can be observed in real-time.

Positive selection can also be detected from cross-population comparisons, based on the prediction that allele frequencies should differ between subpopulations if positive selection has acted in only one of them (Lewontin & Krakauer 1973; Sabeti *et al*. 2007; Akey *et al*. 2010). While such tests do not require time-course data, they remain limited to scenarios where selection acted only in a subset of individuals.

The most broadly applicable strategy for identifying positive selection is to search for its signatures in a single population sample, taken at a single point in time. Approaches from this category aim to identify the characteristic signatures of selective sweeps (Maynard Smith & Haigh 1974; Kaplan *et al*. 1989; Barton 2000), which include a local trough in genetic diversity around the selected locus (Kim & Stephan 2002), characteristic biases in the frequency distributions of single nucleotide polymorphisms (SNPs) (Braverman *et al*. 1995; Fay & Wu 2000), and the presence of a long haplotype that extends much farther than expected under neutrality (Sabeti *et al*. 2002). These signatures form the basis for most popular scans for selective sweeps (Vitti *et al*. 2013).

However, positive selection may not always produce selective sweeps. The classic selective sweep model presupposes that adaptation occurs from a single *de novo* mutation (Hermisson & Pennings 2005). Yet adaptation could often proceed from alleles that are already present as standing genetic variation (SGV) (Orr & Betancourt 2001; Innan & Kim 2004; Barrett & Schluter 2008). This should be particularly common in the evolution of polygenic traits, such as body size, where multiple trait-affecting alleles may be segregating in the population at any time (Pritchard *et al*. 2010).

Whether adaptation from SGV still produced sweep-like signatures depends on the initial frequency and age of a selected allele at the time when positive selection commences (Przeworski *et al*. 2005; Pennings & Hermisson 2006b). If the selected allele has been around long enough to recombine onto different haplotypes prior to the onset of positive selection, several haplotypes may then increase in frequency simultaneously. In this case, diversity is not necessarily reduced in the vicinity of the selected site and SNP frequency spectra can actually become biased towards intermediate frequencies (Przeworski *et al*. 2005). Very similar patterns are produced when adaptation involves several *de novo* mutations that independently emerged on distinct haplotypes, which is expected in very large populations or when mutational target sizes are large (Pennings & Hermisson 2006a; Karasov *et al*. 2010; Messer & Petrov 2013). The patterns generated by adaptation from SGV and recurrent *de novo* mutation are commonly referred to as soft selective sweeps, in contrast to the classical hard selective sweep, where only a single haplotype rises in frequency (Hermisson & Pennings 2005).

Most scans for positive selection have been designed and tested exclusively under the assumption of a hard selective sweep model and we do not know whether they provide a comprehensive picture of the mode and frequency of positive selection, or whether they identify only a subset of instances that is biased towards hard selective sweeps. Simulation studies have shown that selection scans quickly lose power for adaptation from SGV as the initial frequency of the selected allele increases (Przeworski *et al*. 2005; Teshima *et al*. 2006; Garud *et al*. 2015). However, it is unclear whether the alleles involved in adaptation from SGV are typically rare or frequent prior to the onset of selection.

Here we use a set of known quantitative trait loci (QTLs) in the domestic dog (*Canis lupus familiaris*) as positive controls to examine the performance of popular selection scans in a real biological system. Our positive control loci were identified by genome-wide association studies, rather than selection scans, and thus are not necessarily biased towards hard selective sweeps from the outset. We focus specifically on a subset of QTLs for which we know the causal mutations and could thus measure their frequencies in individual dog breeds. This information allowed us to assess which mutations have likely experienced positive selection in which breeds.

There are over 400 dog breeds today that have been bred for highly specific and diverse physical traits, including coat color, size, skull shape, and behavioral traits such as obedience, herding, and hunting. Modern dogs were the first animal to be domesticated, before cattle and horses, and domestication from their wolf ancestors goes back at least 15,000 years. Breeding programs throughout history, however, have resulted in periodic population bottlenecks, inbreeding, high levels of linkage disequilibrium in individual breeds, and a prevalence of inherited diseases such as cancer, heart disease, and hip dysplasia, among others (Lindblad-Toh *et al*. 2005). These features make purebred dogs a particularly challenging system for population genetic analysis.

## RESULTS

### A set of 12 positive controls for studying the signatures of positive selection in dogs

The molecular basis of morphological phenotypes selected during domestication of dog breeds has been extensively studied, and dozens of QTL for breed-specific phenotypes have been identified, which often explain surprisingly high fractions of phenotypic variance (Rimbault *et al*. 2013). We compiled a set of 12 known QTLs distributed across nine chromosomes of the dog genome for which we know the specific mutations that are likely causal for breed-specific traits (Table 1). Our set includes mutations affecting body size (IGF1R, STC2, GHR, and IGF1 (Rimbault *et al*. 2013)), fur type (MC5R (Hayward *et al*., under review) and KRT71 (Cadieu *et al*. 2009)), coat color (MC1R and TYRP1 (Schmutz & Melekhovets 2012)), hair length (FGF5 (Cadieu *et al*. 2009)), lip morphology (CHRNB1 (Baxter *et al*., in preparation)), ear morphology (MSRB3 (Boyko *et al*. 2010)), and snout length (BMP3 (Schoenebeck & Ostrander 2013)). These loci are representative of loci that show evidence of strong selection based on elevated levels of divergence between breeds (Boyko *et al*. 2010; Akey *et al*. 2010; Vaysse *et al*. 2011). Some QTLs known to be associated with breed-specific morphological traits were intentionally excluded from our analysis, because the causal mutations were either not well-tagged by markers in our data set (e.g. the insertion in the 3’UTR of RSPO2 associated with a furnishings phenotype (Cadieu *et al*. 2009)), or the locus was very close to another locus (e.g. the size-related locus HMGA2 (Rimbault *et al*. 2013) that is only 300 kbp away from MSRB3).

**Table 1:**
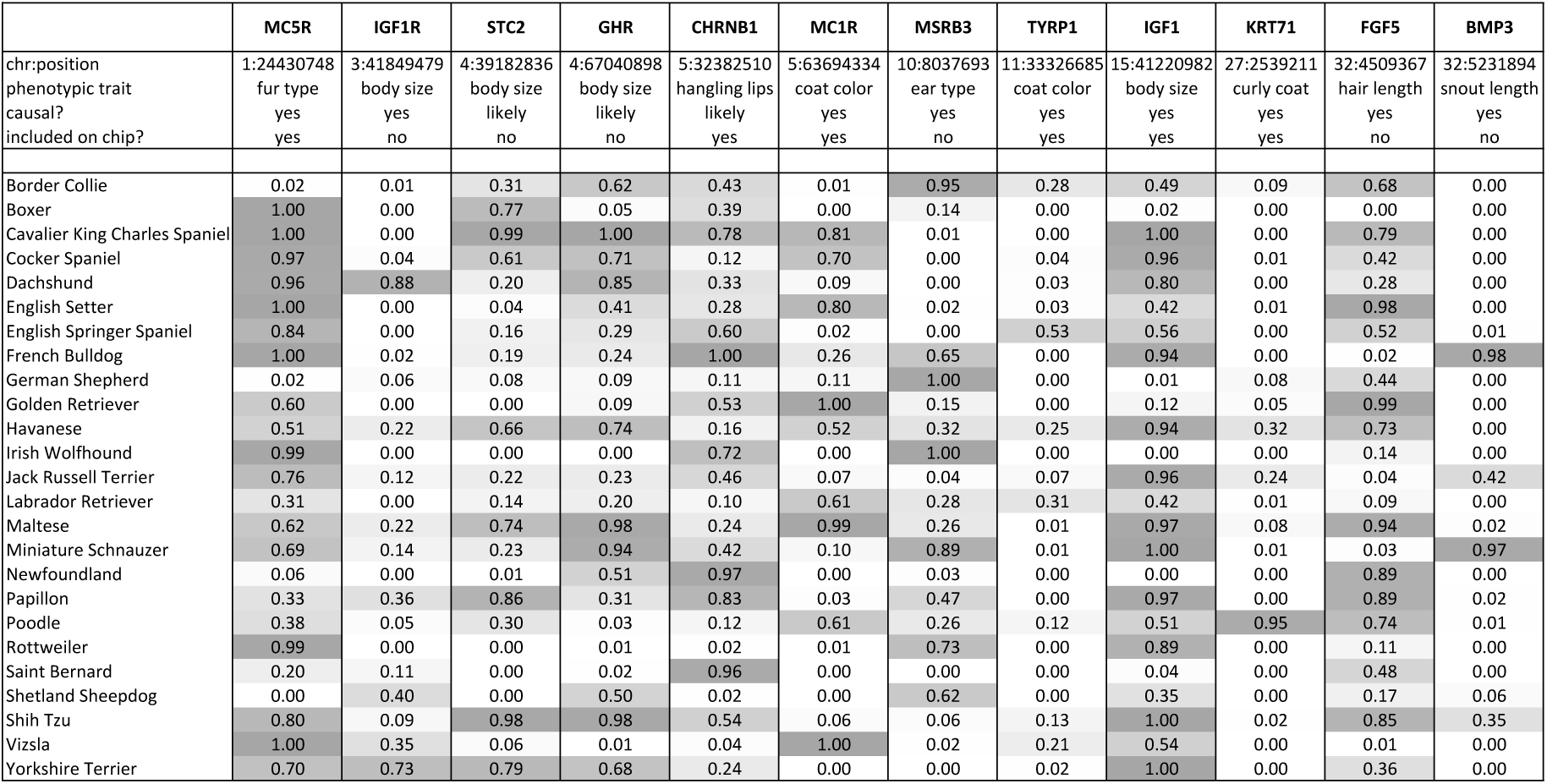
Set of known QTLs with mutation frequencies in individual breeds. The twelve QTLs included in our analysis span a wide range of phenotypic traits that likely experienced positive selection in particular subsets of breeds during the domestication of dogs. For nine of the 12 loci, at least one causal mutation for the phenotypic trait has been identified and for the remaining three loci (STC2, GHR, CHRNB1) we have promising candidate mutations. We focused on one such mutation for each locus (positions are specified in the first row of the table). For six of the 12 loci, these mutations are included on the genotyping chip. We studied 25 dog breeds in our analysis. Numbers in the cells specify the frequency of the known/likely causal mutations in each particular breed (Methods).

We analyzed the population genetic signatures we observed around these 12 loci in population samples from 25 dog breeds, spanning a broad range of morphological variation (Table 1). For each of the 25 breeds, we genotyped a random sample of 25 dogs at ~180,000 SNP markers, using a semi-custom SNP array (Methods). For six of the 12 loci, the known/likely causal mutations are included on the chip. Genotypes were then phased and imputed on the whole set, yielding 50 haploid genomes for each of the 25 breeds (1250 genomes over the whole data set, see Methods). We polarized SNPs using allele information from Culpeo Foxes for SNPs where such information was available (99.46%) and assumed the minor allele to be the derived allele otherwise. To assess whether a particular mutation was likely under positive selection in a particular breed, we estimated population frequencies for the focal mutation at each of the 12 loci in each of the 25 breeds (Table 1, Methods).

### Genome-wide selection scans in 25 dog breeds

We first used the hapFLK statistic (Fariello *et al*. 2013) to confirm that our 12 positive controls indeed show signatures of positive selection in cross-population comparisons. hapFLK was developed to detect differences in haplotype frequencies across many populations, using an F_ST_-based framework that also incorporates information about the hierarchical structure of the populations. Figure 1 shows the results from our genome-wide hapFLK scan including all 25 breeds (only the chromosomes that contain positive controls are shown). Each of our 12 controls is associated with a significant peak (*P*<0.05) in the hapFLK scan, with 7 of the 12 detected as extreme outliers (*P*<0.001). Figure S1 shows the underlying hierarchical breed structure inferred by hapFLK.

**Figure 1:**
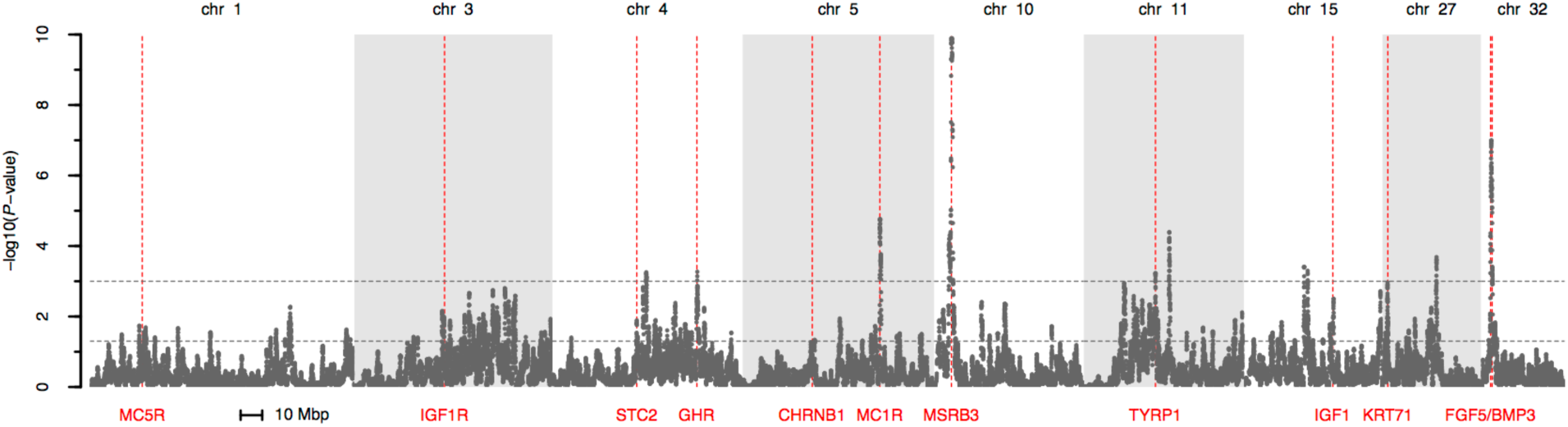
HapFLK results. The figure shows the results from the hapFLK scan performed over all 25 breeds. Results are shown only for those chromosomes that contain at least one of our control loci. The genome-wide thresholds corresponding to *P*<0.05 and *P*<0.001 are shown as horizontal dashed lines. The locations of the control loci are indicated by vertical red lines.

To test whether positive selection at our control loci has also left detectable signatures in the patterns of genetic variation in individual breeds, we ran genome-wide scans using seven popular statistics for identifying sweep signatures from a single population sample. We studied both SNP frequency-based and haplotype-based statistics.

Tajima’s *D* is a popular frequency-based statistics that compares the number of segregating sites (*s*) in a population sample with levels of heterozygosity (*π*) to detect genomic regions with an excess of low or high frequency SNPs compared to neutral expectations (Tajima 1989). Another widely-used statistic is CLR, which underlies the programs Sweepfinder (Nielsen *et al*. 2005) and SweeD (Pavlidis *et al*. 2013). We included both Tajima’s D and CLR as two classic representatives of frequency-based statistics in our study. We also included pairwise heterozygosity per nucleotide (*π*).

Haplotype-based statistics search for elevated levels of haplotype homozygosity expected around a sweep locus. One of the most popular approaches in this category is integrated haplotype score (iHS), which searches for loci where the derived allele resides on a longer haplotype than the ancestral allele (Voight *et al*. 2006). In addition to iHS, we also included the nSL statistic, a recent modification of iHS that has improved power in detecting soft sweeps (Ferrer-Admetlla *et al*. 2014). Note that iHS and nSL are both targeted at the identification of incomplete sweeps, where the selected allele is not fixed in the sample. We further included the H12 statistic that has been developed for the detection of both hard and soft sweeps (Garud *et al*. 2015). Finally, we included a simple haplotype statistic (H) that measures the average length of pairwise haplotype homozygosity tracts around each SNP in base pairs (Methods).

All statistics except H require specification of analysis parameters. For iHS and nSL, minimum haplotype homozygosity levels need to be specified below which haplotypes are no longer extended. In order to improve sensitivity, iHS and nSL are also combined over neighboring data points, which introduces a window-size parameter (Voight *et al*. 2006). CLR requires the specification of the number of grid-points along the chromosome. H12, Tajima’s D, and *π* require specification of the length of an analysis window over which their values are estimated. These windows are typically defined in terms of a fixed number of segregating sites. Given that SNPs in our data were estimated from all 25 breeds, we can either define such windows using all SNP in our data set, or only those SNPs that are actually segregating in the particular breed of interest. We decided to include all SNPs when defining window sizes in order to make results comparable between breeds. Note that this may be considered an “unfair” advantage to the Tajima’s D and H12 statistics, as it incorporates cross-population information: Consider, for example, a window of 25 neighboring SNPs identified using information from all breeds, for which diversity is depleted entirely in a particular breed (*π* = 0). Tajima’s D will then be very negative in this region and H12 will yield a value of one, as only a single haplotype will be present in the window. However, it turns out that in practice the performance of these statistics is not strongly affected by whether we define segregating sites using all 25 breeds, or just the particular breed for which the given statistic is estimated, as we will show below.

Figure 2 shows the results of the seven statistics (iHS, nSL, H, H12, CLR, Tajima’s D, *π*) for the example of French Bulldogs. Different statistics vary markedly in appearance and statistical properties, although some statistics are more similar than others. As expected, iHS and nSL identify largely overlapping candidate regions. Likewise, H12 and H behave similar to each other, consistent with the fact that both statistics measure local levels of haplotype homozygosity (although H12 measures homozygosity over a window of fixed size, whereas H measures the average length of pairwise homozygosity tracts around a SNP in the sample). Tajima’s D and *π* yield similar results as well, suggesting that the signal in Tajima’s D in regions with negative values is driven primarily by local reductions in *π*. Increasing window sizes generally tends to smoothen results for the window-based statistics, reducing noise at the price of lowered sensitivity. Figure S2 shows the results of the scans in French Bulldogs when defining windows using only those SNPs that are actually segregating in our sample. Results are almost indistinguishable between the two approaches, suggesting that our choice of defining window sizes using all SNP in the data set does not have a large effect on the analysis. Results of the selection scans for all 25 breeds are presented in Figure S3.

**Figure 2:**
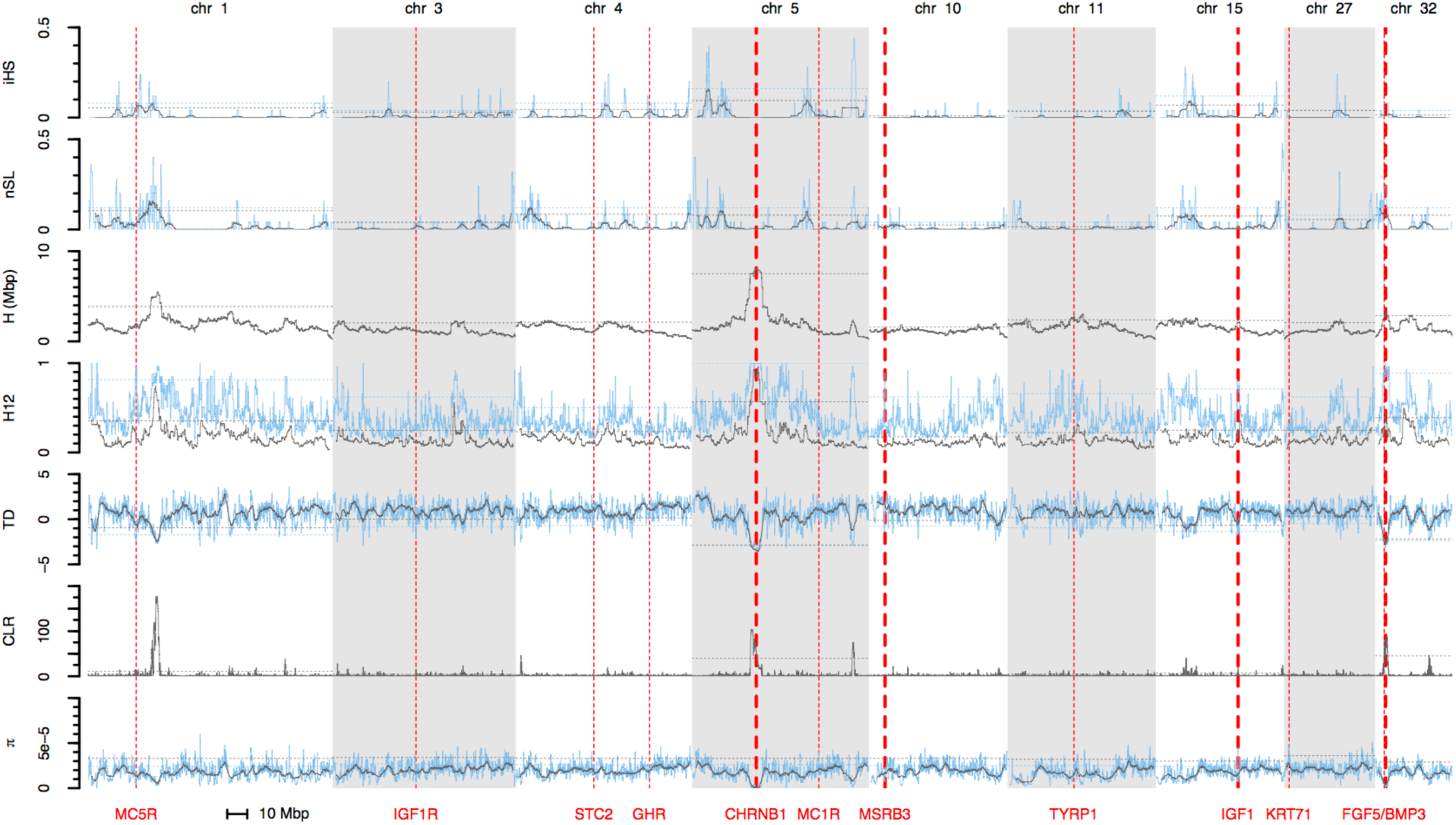
Single-population selection statistics in French Bulldogs. Results for iHS, nSL, H, H12, Tajima’s D, CLR, and *π* along those chromosomes that harbor at least one of our positive controls. For iHS, nSL, H12, Tajima’s D, and *π*, the blue lines show results for a window size of 25 SNPs, grey lines show results for a larger windows of 201 SNPs. Note that signals of positive selection correspond to higher values of iHS, nSL, H, H12 and CLR, but lower values of *π* and more negative values of Tajima’s D. Horizontal dashed lines indicate the 95% quantile cutoffs for the given statistic and window size, which we estimated for each chromosome separately. The positions of the controls are indicated by vertical red lines. The width of these lines corresponds to the frequency at which the causal mutation was observed in the breed in our sample (thin lines: low frequency; thick lines: high frequency). Scans for all 25 breeds are presented in Figure S3.

Note that our SNP data were obtained from a genotyping chip, rather than direct sequencing (Methods). Low-frequency SNPs are therefore underrepresented. This should systematically bias Tajima’s D values towards more positive values and may also affect the CLR statistic. However, since we expect these biases to be present genome-wide, relative comparisons between different regions along the genome should remain informative. Note also that levels of nucleotide and haplotype diversity vary widely between breeds and that our data set covers a wide range of these values (Figure 3).

**Figure 3:**
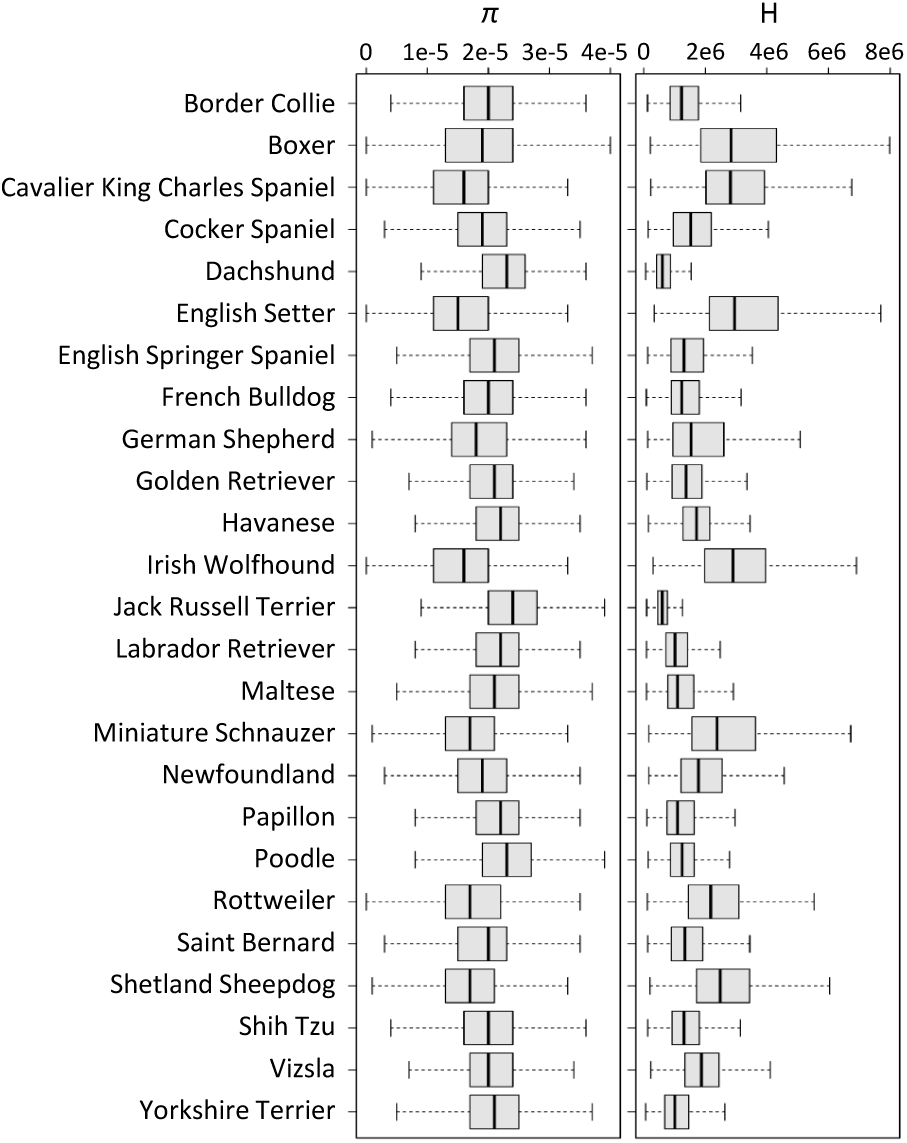
Strong differences in nucleotide and haplotype diversity between breeds. The figure shows the average genome-wide levels of nucleotide heterozygosity (*π*) and length of pairwise haplotype homozygosity tracts (H) in each breed. Values were estimated across all genome-wide SNP positions for the given breed. Values of *π* were estimated using a window size of 51 SNPs. Box plots show medians with first and third quantiles. Note that these values were obtained from our genotyping chip, which comprises only a subset of polymorphic sites. The true diversity levels will be higher and homozygosity stretches will be shorter.

### Genome-wide outlier characteristics

Our genome-wide scans reveal characteristic differences in the number, sizes, and distributions of “peaks” identified by the seven selection statistics. To quantify these differences, we assigned peaks across the genome using an outlier criterion: We considered all data points with value above a given chromosome-wide quantile threshold (*σ*) as candidates for positive selection. For each such data point, we then defined a peak as the window of radius *d* base pairs around its genomic position. Overlapping peaks were combined into a single peak.

We employed a simple outlier approach, rather than using an explicit neutral null model, as such a model would require information about the particular demographic history of each individual breed. Unfortunately we do not generally know much about these demographic histories, except that they can be complicated and differ profoundly between breeds. Our outlier criterion does not require knowledge of demography, but it cannot provide us with information about false discovery rates. However, in our study we focus on assessing the performance of selection scans at known positive controls, which is conceptually different from the discovery of novel targets in that we are not generally worried about the detection of false positives. Instead, we want to study whether scans correctly place the controls among the top signals genome-wide. Our rationale is that our control loci should be located in or near the regions with the strongest signals. The simple outlier approach allows us to draw general conclusions about the number and distribution of such regions identified by each statistic under a given threshold criteria.

Table 2 shows the average number of peaks identified genome-wide per breed and the average fraction of the genome covered by these peaks, using two different quantile thresholds (*σ*=0.95 and *σ*=0.99) and three window radii (*d*=10,50,250 kbp) for each statistic tested. As expected, lower thresholds and larger peak radii both tend to produce more peaks and larger fractions of the genome covered than higher thresholds and smaller radii. Values range from ~600 peaks identified genome-wide per breed by CLR under the 0.95 criterion with *d*=10 kbp, to only ~10 peaks identified genome-wide per breed for iHS under the 0.99 criterion with *d*=250 kbp. Note that nucleotide heterozygosity is very low in our data set: on average, *π*~10^−5^ per site for the breeds in our data set (Figure 3). Thus, neighboring SNPs tend to be several kbp apart, which is why we chose rather large windows.

**Table 2:**
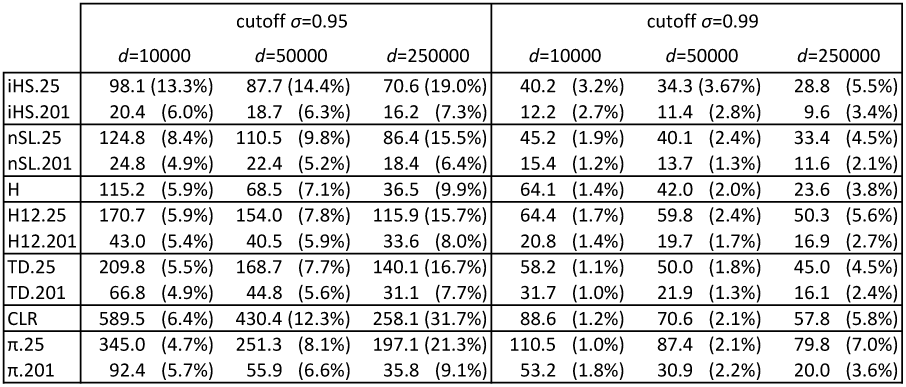
Genome-wide peak statistics. The table shows the number of peaks identified genome-wide by each statistic for a given quantile threshold (*σ*) and window radius (*d*), averaged across all breeds. The numbers in brackets specify the average percentage of the genome that is covered by the peaks in the particular scenario.

### Performance of selection scans at positive controls

We next assessed the performance of each statistics in identifying signals of positive selection at each positive control locus. This was done by measuring the distance between the causal sweep mutation and the next data point with a value above the 0.95 chromosome-wide quantile of the given statistic. If the statistic yielded a value above the 0.95 quantile at the actual causal mutation, we set the distance to zero. We used chromosome-wide quantiles, rather than genome-wide quantiles, because levels of nucleotide and haplotype diversity vary systematically between the different chromosomes within a breed (Figure S3).

A close distance between a causal mutation and an outlier data point is not itself a clear indication that the given statistic has high power in detecting the locus. The close distance could simply be due to chance if values of the statistic fluctuate fast along the chromosome, so that any random genomic position would typically be close to a data point with value above the 0.95 threshold. To assess the significance of a measured closest distance, we therefore calculated empirical *p*-values for observing the given or a shorter distance by chance, based on the distribution of closest distances at random genomic locations in the particular chromosome and breed. Note that these empirical *p*-values are not *p*-values in the regular sense obtained from a neutral null model, but simply indicate the extent to which the observed distance is an empirical outlier regarding the chromosome-wide distribution.

The resulting *p*-values for all locus/breed combinations in which the causal allele has a frequency of at least 50% are shown in Table 3. For the window-based statistics, we show result for window sizes 25, 51, 101, and 201 SNPs. The actual distances between the causal mutation and the closest outlier are provided in Table S1.

**Table 3:**
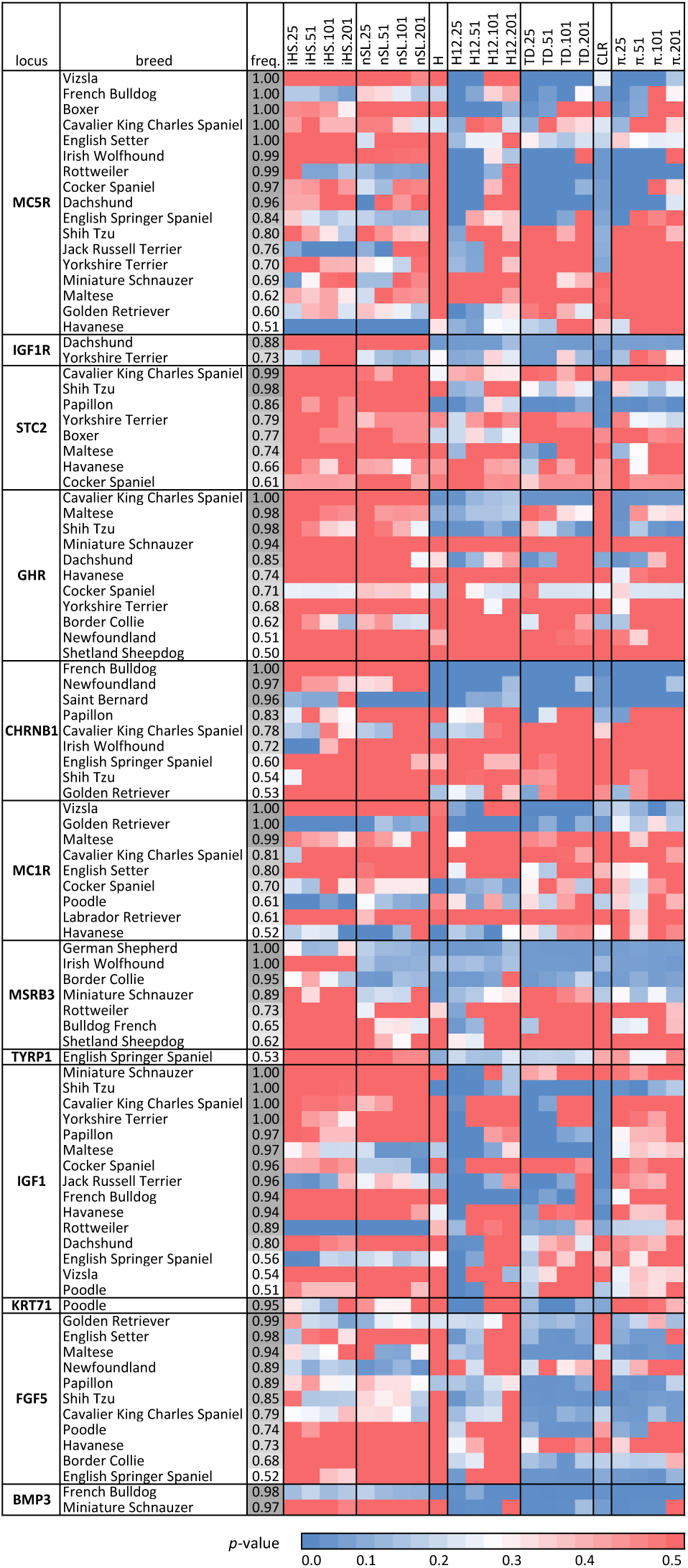
Performance of selection scans at individual QTLs. The table shows for each locus the breeds ordered by the frequency of the causal allele in the particular breed (only breeds with frequency above 50% are shown). The coloring of the cells specifies the *p*-value of the measured distance between the causal mutation and the closest data point that lies above the 95% threshold for the given statistic. Our empirical *p*-values were calculated from the empirical distributions of closest distances for random genetic loci. Different statistics vary widely in whether they detect signatures of positive selection for a given locus/breed combination. In general, signatures of positive selection tend to be detected more frequently, the higher the frequency of the selected mutation in the specific breed.

Table 3 shows that there is substantial variation in the ability to detect signatures of positive selection among different statistics, loci, and breeds. As expected, iHS and nSL produce rather similar results. Interestingly, H12, Tajima’s D, and *π* also appear to be more similar to each other than to the other statistics. H12 and Tajima’s D identify the largest number of locus/breed combinations, at least when using the small windows size of 25 SNPs (Table 4). iHS and nSL identify only one or two (depending on window size) of the 15 fixed sweeps under a 0.05 significance level. They fail to identify any fixed sweep when using a stricter 0.001 significance level. These particular results for iHS and nSL are not surprising, given that both statistics were designed to detect incomplete sweeps. CLR does identify several sweeps under the 0.05 significance level but also does not detect any sweep under the 0.001 significance level. H and *π* have lower performance than H12 and Tajima’s D but better performance than CLR, iHS, and nSL, especially under the stricter 0.001 significance level.

**Table 4:**
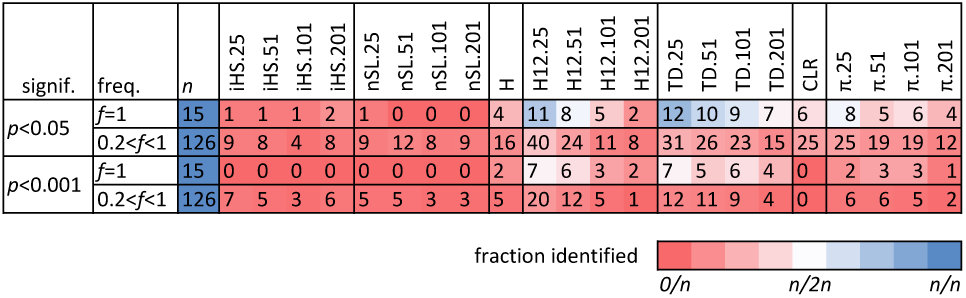
Scan performances under different significance thresholds. The table shows for different sets of locus/breed combinations the number of combinations in which each statistic identifies a signal of positive selection under a significance threshold of *p*<0.05 (top) or *p*<0.001 (bottom). We classified locus/breed combinations into two sets according to whether the selected allele is fixed in our sample (*f*=1) or polymorphic (0.2<*f*<1.0). The “n” column shows the total number of locus/breed combinations in each set. We did not include locus/breed combinations where the selected allele was below 20%.

The similarity between H12, Tajima’s D, and *π* may appear counterintuitive at first glance, given that H12 measures haplotype homozygosity, whereas Tajima’s D and *π* are based on SNP frequencies. A likely reason for this is that we used the number of all SNPs present on our genotyping chip as our estimate for the number of segregating sites (*s*) at a locus. This number is therefore the same for all breeds. However, in those breeds where a sweep has occurred, fewer sites will actually be polymorphic in the window, reducing both *π* and Tajima’s D. At the same time, the number of different haplotypes we expect to observe in the window will decrease as fewer polymorphic sites are present that can break up haplotypes, yielding higher H12 values.

### Haplotype homozygosity levels increase with frequency of selected alleles

Generally we expect that scans should perform better at detecting sweeps the higher the frequency of the selected allele in a particular breed. This tendency is indeed visible in Figure 3 and Table 3. We also observed a clear positive correlation between the frequency of the selected allele in a breed and the value of H at a locus for all loci, except TYRP1 and IGF1 (Figure 4). H simply measures the average haplotype homozygosity lengths among all individuals in the sample. The observation of higher H values for higher-frequency alleles is therefore consistent with the selected alleles residing on longer haplotypes than the ancestral alleles, as more individuals carrying these longer haplotypes will increase the average haplotype lengths among all individuals.

**Figure 4:**
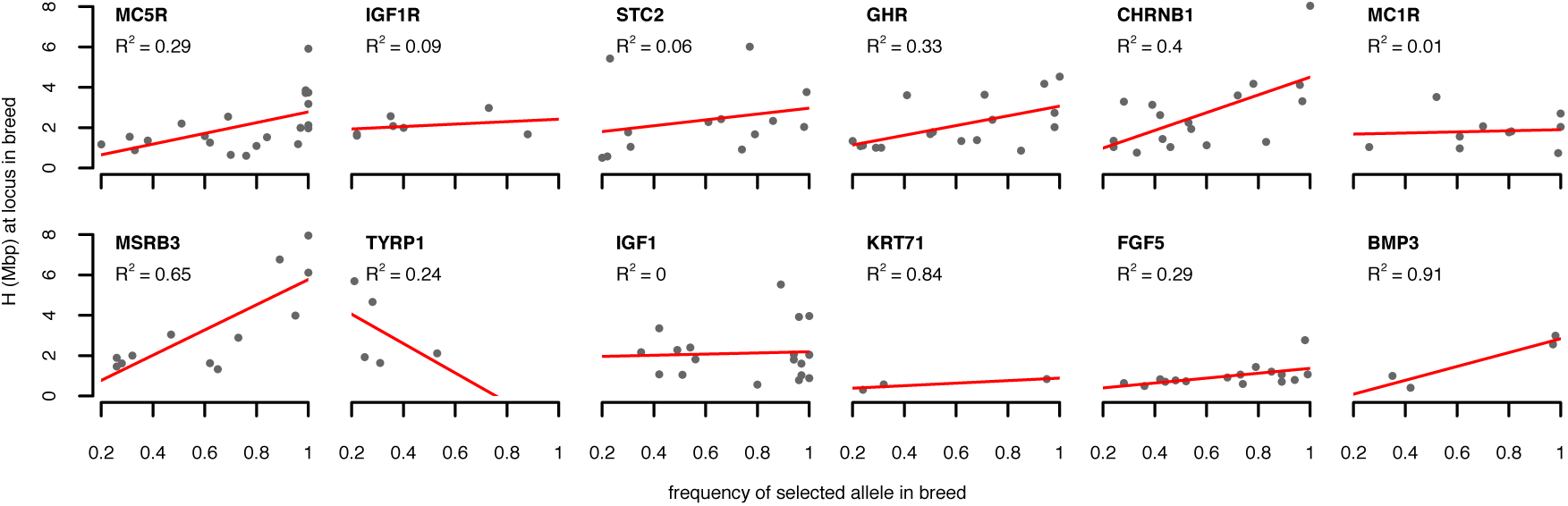
Haplotype homozygosity levels increase with frequency of selected allele. Each panel shows for the particular locus the values of H at the causal site as a function of the frequency in the specific breed (only breeds where the selected allele has a frequency >20% are shown). We observed a positive correlation (measured by R^2^) between allele frequency and the value of H in the breed for all loci except TYRP1 and IGF1.

Note that iHS and nSL lose power to detect a sweep when the selected allele is fixed in the breed (Table 3), as has been observed previously (Schrider *et al*. 2015). As mentioned before, this is expected given that both statistics were specifically designed to detect incomplete sweeps, where both the ancestral and derived allele are still segregating in the population and the haplotypes on which they reside can be compared with each other.

### Positive selection has produced both hard and soft selective sweeps

We analyzed the haplotype patterns and SNP frequency spectra around individual loci in individual breeds to see whether we can understand why some statistics perform better than others at detecting signatures of positive selection in specific cases.

Figure 5a shows the CHRNB1 locus in French Bulldogs, which produced the strongest signal of positive selection in H, H12, Tajima’s D, and *π*. The haplotype and SNP patterns around this locus provide a showcase example of a hard selective sweep. Diversity is depleted over >10 Mbp around the locus (Figure 2). On average, we would expect around 40 sites to be polymorphic over a window of the given size in this breed. However, we do not observe a single polymorphic site at this locus in our sample of 25 French Bulldogs. On average, we would also expect several haplotypes to be present, with the most common haplotype at around 40% frequency. Since no site is polymorphic, we only observe a single haplotype. In contrast to the clear signal identified by H, H12, CLR, Tajima’s D, and *π* at this locus, both iHS and nSL are unable to identify the sweep, consistent with the causal allele being fixed (Figure 2, Table 3).

**Figure 5:**
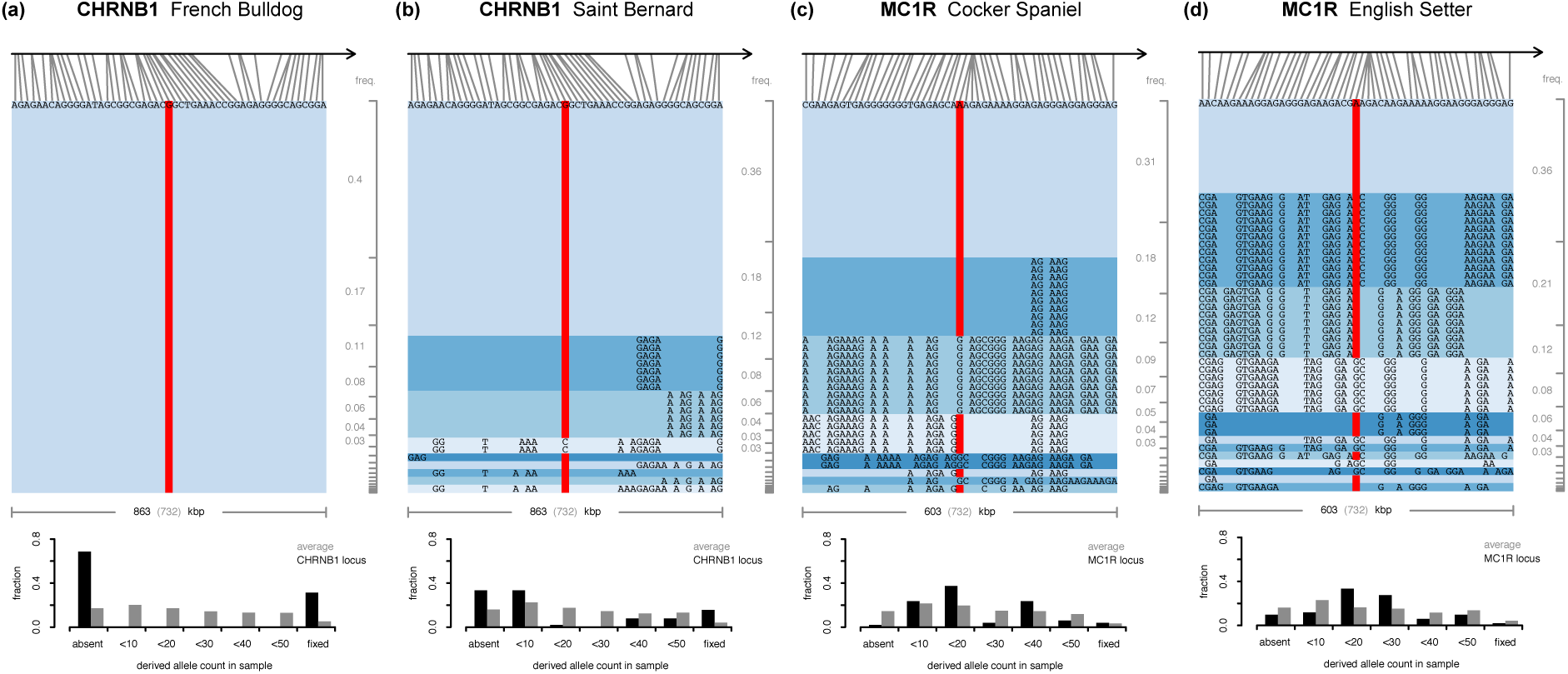
Positive selection produced both hard and soft selective sweeps. Haplotypes and SNP frequency distributions at specific loci in specific breeds. The top part of each panel shows the haplotypes in our sample from the particular breed over a window of 51 sites on our genotyping chip, centered on the causal mutation. The gray brackets on the right show the expected haplotype frequencies ordered by their prevalence in an average window of that size in the chromosome. Bar plots on the bottom of each panel show distributions of SNP frequencies in the window (black), compared with the chromosomal average (gray). The red bars indicate presence of the causal allele. (a) The CHRNB1 locus in French Bulldogs is a hard selective sweep that is fixed in our sample. None of the 51 sites is polymorphic at this locus and only a single haplotype is present. (b) In Saint Bernards, the causal mutation is not fixed in our sample. The most frequent haplotype is at higher-than-expected frequency but several other haplotypes carrying the mutation are also present that may be variants of the major haplotype from recombination and/or mutation events. The SNP frequency spectrum shows the characteristic distortions towards high and low frequencies expected under a hard selective sweeps. (c) At the MC1R locus in Cocker Spaniel the causal mutation is present in 37 of the 50 genomes in our sample. The frequency of the most common haplotype, however, is not much higher than expected by chance and the SNP frequency spectrum is skewed towards intermediate frequencies, compatible with a soft selective sweep. (d) In English Setters, the MC1R locus shows even more pronounced signatures of a soft selective sweep.

In Saint Bernards, for comparison, the mutation at CHRNB1 is at high frequency but two genomes in our sample do not carry it (Figure 5b). Several haplotypes with the causal allele are present at the locus that may be variants of the major haplotype from recombination and/or mutation events early during its sweep (Messer & Neher 2012). The SNP frequency spectrum shows the characteristic distortions of a hard selective sweeps and all scans detect signatures of positive selection at this locus (Table 3).

Figures 5c and 5d show the MC1R locus in Cocker Spaniels and English Setters. Both breeds show signatures strongly suggestive of soft selective sweeps: The frequencies of the most common haplotypes are similar or lower to expectations in an average window, and in both breeds several haplotypes carry the selected mutation. Importantly, some of these haplotypes differ at many sites from each other, including positions right next to the causal site, making it very unlikely that these haplotypes are in fact variants of the same haplotype that arose from mutation or recombination events during the sweep (Messer & Neher 2012). Given that most pure dog breeds are less than 200 years old (Parker 2004; Larson *et al*. 2012), yet some of these haplotype variants are quite common in the sample, it is also unlikely that they arose from recombination events after the sweep. Furthermore, the SNP frequency spectra are atypical for a hard selective sweep as they are skewed towards intermediate frequencies. All of these observations are more consistent with soft selective sweeps where positive selection has driven several haplotypes simultaneously, possibly because selection acted on SGV. Both H12 and H detect signatures of positive selection at MC1R in Cocker Spaniel, other statistics are inconsistent and results strongly depend on window-size. All statistics lack power in identifying signatures of positive selection at this locus in English Setters. Only H12, Tajima’s D and *π* show some signal and only when using short analysis windows (Table 3).

Figure S4 shows haplotype patterns and SNP frequency spectra around the IGF1 locus in 12 different breeds. The selected mutation at this locus has been identified as a SINE element insertion in intron 2 of the IGF1 gene (Rimbault *et al*. 2013) that appears to be absent in Gray Wolves, most large dog breeds, and all wild canids (Gray *et al*. 2010). Hence, we do not expect that positive selection has acted on SGV at this locus, but rather that the selected SINE was a *de novo* mutation that arose during the domestication process. This is largely consistent with the haplotype and SNP frequency pattern in different breeds at this locus, which tend to show signatures of hard selective sweeps.

## DISCUSSION

In our study, we examined the population genomic signatures observed around a set of 12 positive control loci known to have experienced positive selection in specific dog breeds due to their association with desirable morphological phenotypes. The dog system is extraordinary in that it provides a very large number of individual populations (breeds) for which we often know the specific selective pressures experienced. In such a system, the most powerful selection scans should be those that can utilize the information provided by cross-population comparisons, e.g. F_ST_ and XP-EHH based methods (Vitti *et al*. 2013). We confirmed this intuition by showing that hapFLK, a powerful cross-population scan that uses haplotype information in an F_ST_-based framework and incorporates information on the hierarchical structure between breeds, indeed identified all of our controls. However, for many other systems we may not have such cross-population information and will thus rely on scans that can detect signatures of selective sweeps from a single population sample.

Our approach of using positive controls in a real system is conceptually different from previous studies that evaluated the performance of selection scans based on computer simulations (Teshima *et al*. 2006; Huff *et al*. 2010; Poh *et al*. 2014; Lotterhos & Whitlock 2015; Schrider *et al*. 2015). These studies generally assume idealized evolutionary scenarios, such as panmixia, simplified demographic models, and constant parameters over time and space, while interactions between selected sites such as background selection, Hill-Robertson interference, and epistasis tend to be ignored. Unfortunately, we still lack a clear understanding of the importance of these effects and the extent to which they can obscure footprints of positive selection (Bank *et al*. 2014). In addition, many simulation studies assume that adaptation follows the classic selective sweep model. Whether this is an appropriate model for describing adaptation in most biological systems is increasingly being questioned (Pritchard *et al*. 2010; Cutter & Payseur 2013; Messer & Petrov 2013).

We found that artificial selection has indeed left detectable signatures in the polymorphism pattern around our positive controls in purebred dogs. However, whether such signatures were detected varied widely between loci, individual breeds, the particular statistic used, and the choice of analysis parameters. Interestingly, one of the most popular haplotype-based statistics, iHS, proved to be less accurate in identifying signatures of positive selection at our controls than the other statistics, including simpler haplotype-statistics such as H12 and H, as well as the frequency-based statistics CLR, Tajima’s D, and *π*. This could be due to a number of reasons: It is well known that iHS has difficulties identifying fixed sweeps because it requires the ancestral allele to be segregating in the population (Schrider *et al*. 2015). We indeed observed that both iHS and nSL had particularly low power at those locus/breed combinations where the causal allele was fixed in our sample (Table 4). Furthermore, the generally high levels of LD in purebred dogs (Sutter *et al*. 2004; Lindblad-Toh *et al*. 2005; Boyko *et al*. 2010) could limit the sensitivity of haplotype-based statistics, as only extremely strong sweeps may be able to generate haplotypes that are even longer than those already present. Note, however, that two other haplotype-based statistics, H and H12, identified many positive controls.

The H12 statistic estimated over short windows of 25 segregating sites identified the largest number of positive controls in our study, followed by Tajima’s D and *π*. This finding suggests that the signals of positive selection identified by these three statistics may be largely driven by the difference between the local density of SNPs on our genotyping chip (which we used for calculating the number of segregating sites in Tajima’s D as well as for defining the window length for estimation of H12 and *π*) and the number of SNPs that are actually polymorphic in a particular breed in the given window.

Purebred dogs are clearly an exceptional system, characterized by strong artificial selection that is sometimes even repeatable between breeds (Boyko *et al*. 2010). In addition, phenotypic variance for breed-defining morphological traits is often explained by surprisingly few mutations (Rimbault *et al*. 2013). As such, purebred dogs provide an excellent system for mapping the genetic basis of positively selected variants.

However, some aspects of our data set could confound the results in our study. First, because SNPs were obtained from a genotyping chip, rather than direct sequencing, they should be biased towards common variants, which might compromise the performance of frequency-based methods such as CLR and Tajima’s D. In addition, the high levels of LD in dogs due to increased inbreeding could limit the power of haplotype-based methods. Dog breeds also vary in effective population size by several orders of magnitude (Leroy *et al*. 2013), overlapping the range observed in smaller natural populations. In many ways, detection of selective sweeps in smaller populations is more difficult than in large populations as extensive drift can obscure and weaken the signatures of sweeps.

The severe bottlenecks during the breeding process could have systematically affected the patterns generated by positive selection, such as whether hard or soft sweeps should be more common. For example, recurring bottlenecks can have “hardened” sweeps from SGV that were initially soft (Wilson *et al*. 2014). The mode and signatures of adaptation in large natural populations may therefore be quite different from those observed in purebred dogs and additional work is needed to evaluate the performance of methods for detecting selective sweeps in such populations.

## METHODS

### Genotyping

Genotyping data are from Hayward *et al*. (in review). Briefly, blood was collected through cephalic venipuncture under Cornell IACUC # 2005-0151, and genomic DNA was extracted using a standard salt precipitation from EDTA blood samples and stored in the Cornell Veterinary Biobank.

Genotyping was done using the Illumina 170k CanineHD array, which was developed using the dog reference sequences (generated from a Boxer and a Poodle) and pooled DNA from a series of European and Asian breeds (Irish Wolfhounds, West Highland White Terriers, Belgian Shepherds, and Shar-Peis) as well as pooled wolf DNA as described in (Vaysse *et al*. 2011). We customized this array by adding 12,143 markers ascertained from whole genome sequencing data from mostly Eurasian village dogs (Auton *et al*. 2013), approximately equally split between East Asian and Western dogs. Markers were preferentially chosen for being in coding regions but poorly tagged by existing array markers. The genotypes were combined with published CanineHD data from (Axelsson *et al*. 2013). The full SNP panels (3 million SNPs for the CanineHD array design and 14 million SNPs for the custom array content) was pruned for evenness, ability to design probe sequence, and efficiency. In general, no effort was made to differentially enrich one source or another in particular regions of the genome, except that a subset of custom SNPs were specifically included in the IGF1 and MSRB3 regions to facilitate fine-mapping of those loci. No such enrichment of markers was made for the other 10 loci.

The un-imputed dataset contained a call rate over 99.1%, and no locus contained >5% missing data. Imputation was done because some methods to detect positive selection require no missing data, but the proportion of imputed genotypes is negligible and unlikely to bias the results.

Phasing was done for all autosomal and X chromosome markers with minor allele frequency (MAF) > 0.01 using SHAPEIT (Delaneau *et al*. 2013). Select regions showing strong evidence of positive selection when comparing allele frequency data across breeds and associated with a known phenotypic effect were chosen for analyzing selection signatures in each population.

### Frequency estimates of causal mutations in breeds

Selection signatures were estimated from a randomly selected subset of 25 unrelated individuals per breed. The allele frequency of the causal variant (when known) or the top associated variant was estimated from the entire dataset (Hayward *et al*. in review) based on a much larger number of individuals genotyped (25 to 722 dogs per breed).

### Selection scans

The hapFLK statistic was calculated using the program hapflk (version 1.2) (Fariello *et al*. 2013), downloaded from: https://forge-dga.jouy.inra.fr/projects/hapflk (August 2015). The population tree was obtained by hapFLK to compute Reynolds distances and the kinship matrix across all 25 breeds genome-wide, using Culpeo Fox as the outgroup. The hapFLK scan was run using all 25 breeds genome-wide. We used the following parameters: 8 clusters (-K 8), 20 EM runs to fit the LD model (-nfit=20), phased data (--phased). Once hapFLK values were generated, we calculated P-values by fitting a standard normal distribution genome-wide in R (Fariello *et al*. 2013).

iHS scans were performed using the program selscan (version 1.0.4) (Szpiech & Hernandez 2014), downloaded from: http://github.com/szpiech/selscan (April 2015). All scans were run on polarized data with default iHS selscan parameters: --max-extend 1000000 (maximum EHH extension in bp), --max-gap 200000 (maximum gap allowed between two SNPs in bp), --cutoff 0.05 (EHH decay cutoff). We used the recombination map of Auton *et al*. (Auton *et al*. 2013). The output results for each SNP were then frequency-normalized over all chromosomes using the script norm, provided with selscan. This normalization was also done using default parameters: --bins 100 (number of frequency bins). The fractions of SNPs with values above 2.0 were calculated over genomic windows of specified sizes (25, 51, 101, 201 neighboring sites on our chip) and the resulting ratio was assigned to the position of the center SNP of the window, as suggested in (Voight *et al*. 2006).

In contrast to iHS, which measures the length of haplotypes in terms of genetic distance and thereby requires specification of a recombination map, the nSL statistic measures haplotype lengths in terms of the number of segregating sites in the sample, making it more robust to recombination rate variations. nSL scans were performed using the original implementation of the statistic (Ferrer-Admetlla *et al*. 2014), downloaded from: http://cteg.berkeley.edu/~nielsen/resources/software/ (April 2015). All scans were run using default nSL parameters. The output results were normalized and averaged over windows following the same procedures used for iHS.

The H statistic was estimated using the program H-scan (version 1.3), downloaded from: http://messerlab.org/software/ (April 2015). The H statistic measures the average length of pairwise haplotype homozygosity tracts around a given genomic position in base pairs. The length of the homozygosity tract *h_ij_*(*x*) for a pair of samples (*i,j*) at genomic position *x* is defined as the distance between the first heterozygous site to the left and to the right of *x*. The value of *H*(*x*) at position *x* is then defined as the average over all pairs in the sample: *H*(*x*) = 2/(*n*(*n*-1)) Σ*_i_*_<_*_j_ h_ij_*(*x*). H values were calculated at each SNP position in the data set. All scans were run using default H-scan parameters.

H12, Tajima’s D, and *π* values were calculated over windows of a fixed number of segregating sites (*d*), 25, 51, 101, and 201, defined by the number of SNPs on our genotyping chip. The values of each statistic estimated over a window were assigned to the position of the center SNP of that window. H12 values were estimated following the definition provided in (Garud *et al*. 2015). Tajima’s D values were variance-normalized according to the formulas given in (Tajima 1989). Note that because all scans were run on a fully imputed data set, haplotype clustering for H12 is unambiguous in this study.

CLR is a likelihood-ratio test that compares the SNP frequency spectrum in candidate regions with the genomic background in order to identify regions with sweep-characteristic deviations. CLR scans were performed using the software SweeD (version 3.1) (Pavlidis *et al*. 2013), downloaded from: http://sco.h-its.org/exelixis/web/software/sweed/ (April 2015). For each chromosome CLR was calculated with a resolution of 10000 bins, assuring that the density of bins is much higher than the density of SNPs in each chromosome. All CLR scans were run on unfolded spectra using the polarized data.

## ACKNOWLEDGMENTS

The authors thank Juan Felipe Beltran for programming support and three anonymous reviewers for comments and suggestions. FS was supported by a Presidential Life Science Fellowship from Cornell University. PWM was supported by startup funds from Cornell University. ARB was supported by the National Institute of General Medical Sciences and the National Institute of Aging of the National Institutes of Health under awards 12-PAF-04410 and AG044284-01.

## DATA ACCESSABILITY

All data will be deposited upon acceptance of the manuscript at Dryad (datadryad.org).

